# Rescue of *Mycobacterium bovis* DNA obtained from cultured strains during official surveillance of animal TB: key steps for robust whole genome sequence data generation

**DOI:** 10.1101/2023.08.24.554648

**Authors:** Daniela Pinto, Gonçalo Themudo, André C. Pereira, Ana Botelho, Mónica V. Cunha

**Affiliations:** Centre for Ecology, Evolution and Environmental Changes (cE3c) & CHANGE - Global Change and Sustainability Institute, Faculdade de Ciências, Universidade de Lisboa, Lisboa, Portugal; Biosystems & Integrative Sciences Institute (BioISI), Faculdade de Ciências, Universidade de Lisboa, Lisboa, Portugal; National Institute for Agrarian and Veterinary Research (INIAV IP), Av. da República, Quinta do Marquês, 2780-157 Oeiras, Portugal

**Author notes:** Correspondence: Mónica V. Cunha, Centre for Ecology, Evolution and Environmental Changes (cE3c), Faculdade de Ciências, Universidade de Lisboa, Campo Grande, 1749-016 Lisboa, Portugal.

**Keywords:** animal tuberculosis, *Mycobacterium bovis*, whole genome sequencing, whole genome amplification, computational biology, mixed infection

## Abstract

Epidemiological surveillance of animal tuberculosis (TB) based on whole genome sequencing (WGS) of *Mycobacterium bovis* has recently gained track due to its high resolution to identify infection sources, characterize the pathogen population structure, and for contact tracing. However, the workflow from bacterial isolation to sequence data analyses has several technical challenges that may severely impact the power to understand the epidemiological scenario and inform outbreak response.

While trying to use archived DNA from cultured strains obtained during routine official surveillance of animal TB in Portugal, we struggled against three major challenges: the low amount of *M. bovis* DNA obtained from routinely processed animal samples; the lack of purity of *M. bovis* DNA, *i.e.* high levels of contamination with DNA from other organisms; and the co-occurrence of more than one *M. bovis* strain per sample (within-host mixed infection). The loss of an isolate’s genome is a missed link in transmission chain reconstruction, hampering the biological and epidemiological interpretation of data as a whole.

Upon identification of these challenges, we implemented an integrated solution framework, based on whole genome amplification and a dedicated computational pipeline, to minimize their effects and recover as many genomes as possible. With the approaches described herein, we were able to recover 62 out of 100 samples that would have otherwise been lost. Based on these results, we discuss adjustments that should be made in official and research laboratories to facilitate sequential implementation of bacteriological culture, PCR, downstream genomics and computational-based methods. All of this in a time frame supporting data-driven intervention.

## Introduction

*Mycobacterium bovis* is the main causative agent of animal tuberculosis (TB) [1]. This infectious disease of cattle causes significant costs to the livestock sector, as applicable European legislation determines infected animals must be slaughtered and affected herds quarantined [2]. This hampers commercialization of animal-derived products and animal movement. In Portugal, the animal TB eradication program involves passive surveillance at abbatoirs and determines as well that livestock (*i.e.* cattle) must be regularly tested by the single-intradermal cervical comparative test or by interferon-gamma blood testing. However, *M. bovis* infection is not limited to cattle (*Bos taurus*), affecting other livestock species and occurring at the wildlife-livestock interface [3–5]. In the so-called Portuguese TB- epidemiological risk area, where hunting activities are seasonly regular, big game (wild boar, *Sus scrofa*; red deer, *Cervus elaphus*; fallow deer, *Dama dama*) must be preliminarily examined in the field by certified veterinarians. Samples from TB-compatible lesions found in hunted animals or slaughtered cattle are sent to the national reference laboratory, where they are processed by gold standard methods, such as bacteriological culture, followed by molecular identification and differentiation (by PCR) of *Mycobacterium tuberculosis* complex (MTBC) members [6,7].

Methods for culture-based detection of mycobacteria have evolved over the years. Classically, the method includes tissue maceration, decontamination, inoculation into selective solid media, and visual inspection of bacterial colonies. Automatic growth detection systems are now available, speeding up bacterial growth detection and increasing the ability to manage a large number of samples. While based on the classic culture method, the automatic systems rely on an oxygen-sensitive fluorescent dye for bacterial growth detection.

Molecular methods, based on the detection of specific genome targets, have been implemented over the years, either to classify the MTBC ecovar after mycobacterial isolation or to speed up laboratorial diagnosis via the direct detection of *M. bovis* in animal tissue samples [8,9]. These methods range from more traditional protocols, including restriction fragment length polymorphism analysis of the *gyrB* gene [10] and commercially available reverse hybridization assays, to real-time PCR detection of IS*6110*, *mpb70*, or the Regions of Difference (RD) [8].

Over the last years, epidemiological surveillance programs based on whole genome sequencing (WGS) have been gaining track in low prevalence and endemic settings, as shown by the large number of studies performed in France [11], Mexico [12], Portugal [13], Spain [14], Germany [15], USA [16], New Zealand [17], the United Kingdom [18], and Uruguay [19]. Genomic surveillance has major benefits compared to traditional molecular genotyping schemes, such as spoligotyping or mycobacterial interspersed repetitive units-variable number tandem repeats (MIRU-VNTR), due to superior resolution to identify infection sources, characterize *M. bovis* population structure, trace contacts, and inform biosecurity intervention [12]. In developed countries with ongoing official control and surveillance plans, genomic surveillance is being progressively integrated by reference laboratories into routine analyses. This implies the integration in real-time of culture-dependent and molecular methods with next-generation sequencing. Nonetheless, this implementation may raise methodological challenges.

Here, we share our experience as end-users of biological material (*i.e.* archived DNA) obtained from TB official surveillance and routine gold-standard processing (culture, PCR) of tissue samples from animals presumptively infected with *M. bovis*. We used the DNA extracted either from isolated colonies on solid media or from bacterial suspensions grown in automatic detection systems to perform WGS and sequence analyses up to variant [single nucleotide polymorphism (SNP)] calling and *de novo* assembly. We have found three main issues: (i) routine recovery of low amount of DNA, not sufficient for WGS application; (ii) high contamination rate with DNA from other organisms, which compromised the amount and quality of the sequence data obtained; and (iii) samples containing more than one *M. bovis* strains (mixed infection), which precluded correct SNP calling. To overcome each one of these issues, we implemented rescue approaches and recovered a large number of samples that would otherwise be unusable for whole genome-based epidemiological surveillance. This has enabled the recovery of samples and the associated data, which is already of key importance, but has also uncovered steps of the upstream procedure before WGS that could be adjusted to minimize the issues found and further facilitate the integration of culture-dependent *M. bovis* detection with whole genome sequence analyses.

## Materials and Methods

### Sample collection and processing

Tissue samples obtained between 2002 and 2021 from hunted wild boar and red deer, and slaughtered cattle, positive in the intradermal tuberculin test and/or exhibiting suspect TB-lesions during veterinarian inspection, were submitted to the National Reference Laboratory for animal TB (INIAV, I.P.) for analysis.

Sample processing was performed according to the OIE Manual of Diagnostic Tests and Vaccines for Terrestrial Animals [20]. This was done before 2018 by inoculation in Löwenstein–Jensen solid medium and simultaneously with a BD BACTEC™ 9000 automatic detection system (Becton, Dickinson and Company, New Jersey, USA), and from 2019 onwards with a BD BACTEC™ MGIT™ 960 system.

Collected tissues were homogenized with a pestle and mortar, followed by decontamination with 4% sodium hydroxide for 30 minutes (15 minutes from 2016 onwards) to inactivate other contaminating bacteria. After neutralization with hydrogen chloride and centrifugation of the resulting suspension, the sediment was inoculated into Löwenstein– Jensen with pyruvate slants. The inoculated tubes were then incubated at 37°C for up to 42 days, while continuously monitoring the medium for the presence of bacterial colonies. Colonies from positive cultures were visually inspected for the characteristic morphology of *M. bovis*. Some of those isolates of Batch 2 (n = 29) were subcultured in Middlebrook 7H9 (Difco) and passaged in Middlebrook 7H10 (Difco). Both media were supplemented with 5% sodium pyruvate and 10% ADS (50 g albumin, 20 g glucose, 8.5 g sodium chloride in 1 L water) enrichment. Incubation followed at 37°C in a level 3 biosecurity facility.

In parallel, and according to the BD BACTEC™ manufacturer instructions, collected tissues were homogenized and decontaminated as described above, and incubated in Middlebrook 7H9 broth supplemented with OADC and PANTA (Polymyxin B, Amphotericin B, Nalidixic Acid, Trimethoprim, and Azlocillin) and an oxygen-sensitive fluorescent compound. The inoculated tubes were incubated at 37°C for up to 42 days, while continuously monitoring the fluorescence signal representative of bacterial growth. Once the fluorescence crossed the threshold, the sample was deemed positive and readily processed.

Cell biomass (one colony) grown on solid medium or 1 mL of liquid culture suspension were collected, and the cells were recovered by centrifugation. The cell pellet was then washed with 1 mL of phosphate buffer saline (PBS), resuspended in 200 μl of PBS, and heat-inactivated at 99˚C in a water bath for 30 minutes. The suspension was cleared by centrifugation and the supernatant was used for downstream analysis.

### Mycobacterium bovis identification

The presence of *M. bovis* in the whole cell extracts was confirmed by one of three methods: (i) by restriction fragment length polymorphism analysis of the PCR-amplified *gyrB* gene after hydrolysis with *Rsa*I and *SacI*I as described before [10]; (ii) by reverse hybridization assays INNO-LiPA Mycobacteria (Innogenetics, Belgium) or GenoType *Mycobacterium* (Hain diagnostics, Germany), following the manufacturer’s instructions; (iii) by real-time PCR detection of Regions of Difference (RD) 1, 4, and 9 as described elsewhere [8].

### DNA concentration determination

DNA quantification was performed using the Qubit dsDNA HS assay kit or the Qubit dsDNA BR Assay Kit (Thermo Fisher Scientific, Massachusetts, USA) according to the manufacturer’s instructions. This kit is highly selective for double-stranded (ds) DNA and hence ideal for the quantification of dsDNA in crude extracts.

### Whole genome amplification (WGA)

WGA was performed using GenomiPhi^TM^ V2 DNA Amplification Kit (Cytiva, Massachusetts, USA) according to the manufacturer’s instructions from a starting dsDNA amount of up to 10 ng.

### Whole genome sequencing (WGS) and data analysis

The WGS of the samples described in Reis *et al*. [13] (Batch 1) was performed by MiSeq and HiSeq Illumina technology (paired-end 250 pb) at the United States Department of Agriculture (USDA, Ames, IA, USA) with Nextera XT DNA Library Prep Kit from Illumina. WGS of the samples in Batch 2 was commercially conducted by STABVida (Portugal) using Illumina NovaSeq platform with TrueSeq Library Prep kit (paired-end 150 bp). WGS of the samples of Batch 3 was commercially performed by Eurofins (Germany) using Genome Sequencer Illumina NovaSeq platform (Illumina, California, USA) (paired-end 150 bp reads).

The new raw data (Table S1) have been deposited in a public domain server at the NCBI SRA database, under BioProject accession number PRJNA946560.

Upon receiving the read files, a quality control check was performed with a local installation of FastQC (version 0.11.9) [21]. For read filtering, trimming and sequence overrepresentation analysis, FastP (version 0.23.2) [22], hosted at Galaxy server [23], was used with enabled “overrepresented sequence analysis”. The cleaned-up reads were submitted to Kraken2 (version 2.1.1) [24], also hosted at Galaxy server [23], for read taxonomical classification using its Standard database. KrakenTools (https://github.com/jenniferlu717/KrakenTools) were then used to extract reads classified as belonging to the *Mycobacterium* taxon (NCBI tax ID 1763) everytime the *Mycobacterium* read content in the sample was below 70%. Mycobacterial reads were aligned to the genome of *M. bovis* AF2122/97 (NC_002945.4) using the vSNP pipeline (https://github.com/USDA-VS/vSNP). Quality control of the alignments was performed with QualiMap BamQC (version 2.2.2c+galaxy1) [25,26], hosted at Galaxy Europe with 4350 bins, and SNP density per kilobase was determined with VCFtools (version 0.1.16) [27].

### Strain separation

Whenever appropriate, and upon the alert from vSNP of mixed reads, strain separation was performed with SplitStrains [28] starting from the vSNP generated alignment to reference BAM file. The reference was changed to *M. bovis* AF2122/97 (NC_002945.4). The same parameters were used in all the strains: -*s*, specifying the start position on the genome, was set to 1; -*e*, specifying the end position on the genome, was set to 4,349,100; - *fd* was set to 0 to force SplitStrains to not ignore sites based on the depth of coverage (as vSNP also does not); -*u* was set to 99, to ignore proportions only above this value; -*l* was set to 1, to ignore proportions only below this value; -*m* was set to 5, to exclude reads below 5 map quality; -*q* was set to 5, to exclude reads below 5 quality; -*ft* was set to 0 so as to not exclude any position from the analysis.

Upon read separation, reads were extracted from the original BAM alignment files with SplitStrains package script rmreads.py. Each read set was once again aligned to the reference genome with the vSNP pipeline.

### De novo genome assembly

Genomes were *de novo* assembled with the Unicycler pipeline v.0.5.0, available at https://github.com/rrwick/Unicycler [29]. For short read only data, Unicycler assembles genomes *de novo* optimizing the use of SPAdes [30] by evaluating which k-mer size produces the best assembly. We applied a conservative bridging mode to avoid misassembly and the k-mer size was searched and selected between 20-95% of read length. Contigs smaller than 300 bp were removed, considering the reads’ average size, and following SPAdes guidelines [31].

IDBA-UD assembler [32] hosted in Galaxy Europe server was used to generate the assemblies from the samples subjected to WGA. The assembler was run with a minimum k value of 5, a maximum k value of 315 and an increment of k-mer of 10 in each iteration. The seed k-mer size for alignment was set to 15 and minimum size of contig considered was 500 bps. Minimap2 [33] was used for mapping the contigs to the reference genome and Samtools [34] for contig fusion.

To assess the quality of the *de novo* assemblies, we ran the Quast pipeline (http://quast.sourceforge.net/quast.html), which remaps contigs to the *M. bovis* AF2122/97 reference genome, and calculates a number of metrics, such as genome fraction covered [35].

## Results

### Sample collection and M. bovis isolation

As a result of the implementation of the National Control and Eradication Program for Animal Tuberculosis, in Portugal, coordinated by the National Veterinarian Authority (DGAV), the laboratorial confirmation of suspect cases is under the responsibility of the National Reference Laboratory (NRL) for Animal Tuberculosis, INIAV IP. Over the years, the NRL has archived DNA extracted from *M. bovis* isolates obtained during bacteriological culture of cattle or big game tissues.

For this study, we selected a set of archived DNA based on: (i) geographical location and collection year of animal samples, selecting those that were collected in the Epidemiological Risk Area of Animal Tuberculosis for big game species, *i.e.*, near the Spanish border in the Central area of mainland Portugal; (ii) a balanced representation of the main animal hosts of *M. bovis* in Portugal (see below).

We selected 302 samples from the 2002 - 2021 period, in which the presence of *M. bovis* was confirmed by culture-and PCR-dependent methods (Figure 1). Of those, 102 samples originated from cattle (*Bos taurus*), 100 from wild boar (*Sus scrofa*), and 100 from red deer (*Cervus elaphus*). A total of 219 DNA samples were recovered from bacteria grown in solid medium (2002-2018), while 83 were extracted directly from the BD BACTEC™ MGIT™ 960 automatic detection system liquid cultures (2019-2021). Of these, 44 samples were first described in the study of Reis *et al*. [13] (Table S1).

**Figure 1.**
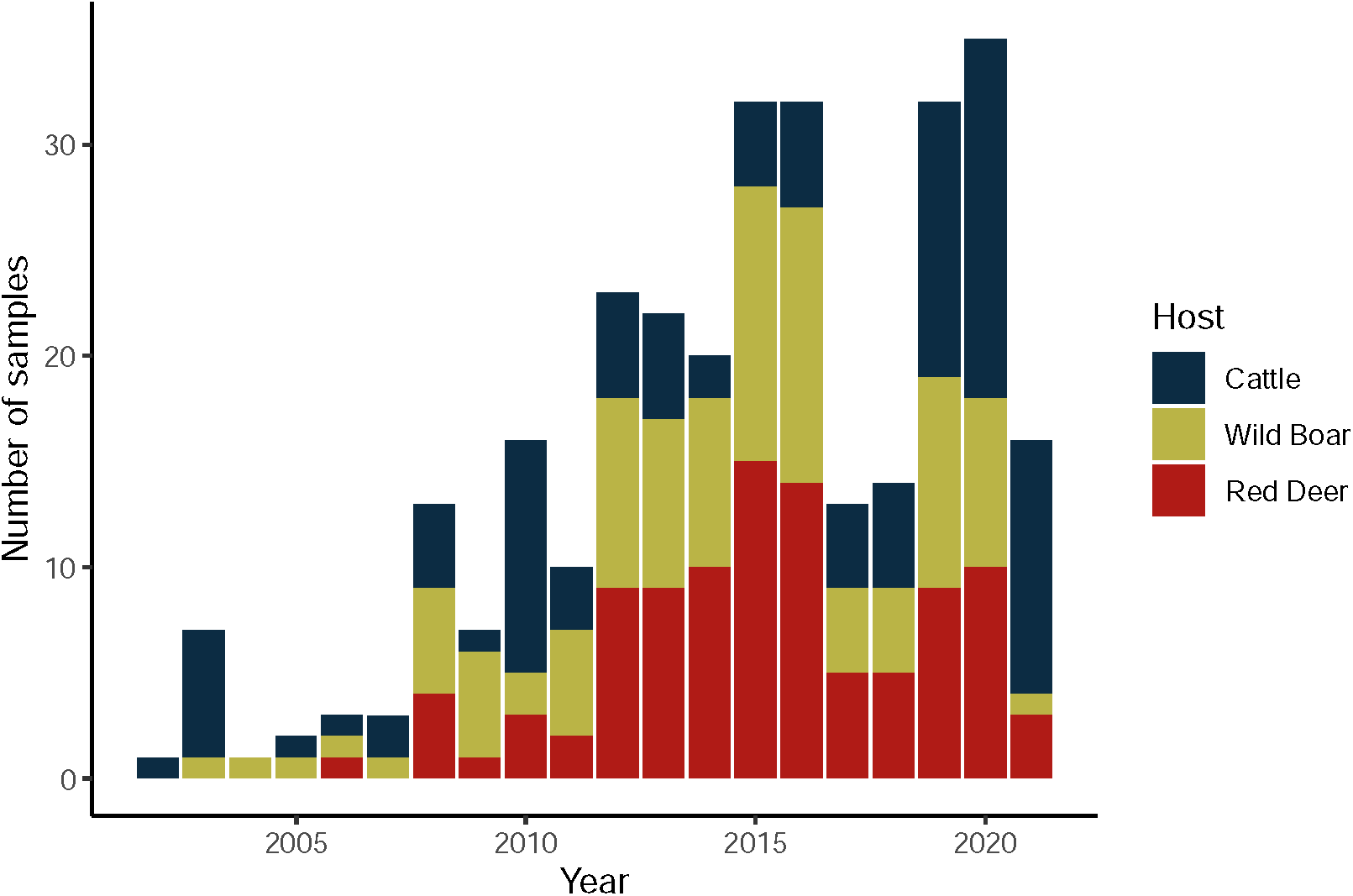
*Mycobacterium bovis* dataset. Distribution of analyzed *M. bovis* samples by year of collection and animal host.

### Recovery of samples containing a low amount of DNA for WGS using WGA

With the aim of including these *M. bovis* isolates in our ongoing whole genome-based epidemiological surveillance, we proceeded with the quantification of the DNA present in the archived whole cell (*i.e.* crude) extracts. We used a highly selective and sensitive method for double-stranded (ds) DNA ideal for quantification in crude extracts. While for the 219 isolates, obtained between 2002 and 2018, the amount of DNA was always enough to proceed with commercial WGS, the same was not true for the most recent samples, obtained between 2019 and 2021. For these, we have detected between 0.1 and 242.9 ng/μl of dsDNA in crude extracts, with several samples containing very low DNA concentrations (Figure 2A). This translated into 250 of the 302 samples (82.8%) having enough DNA to allow direct WGS, being those 100% of the samples before 2018 and only 37.3% of the more recent samples. Hence for these, we proceeded with WGS using paired-end Illumina sequencing technology.

**Figure 2.**
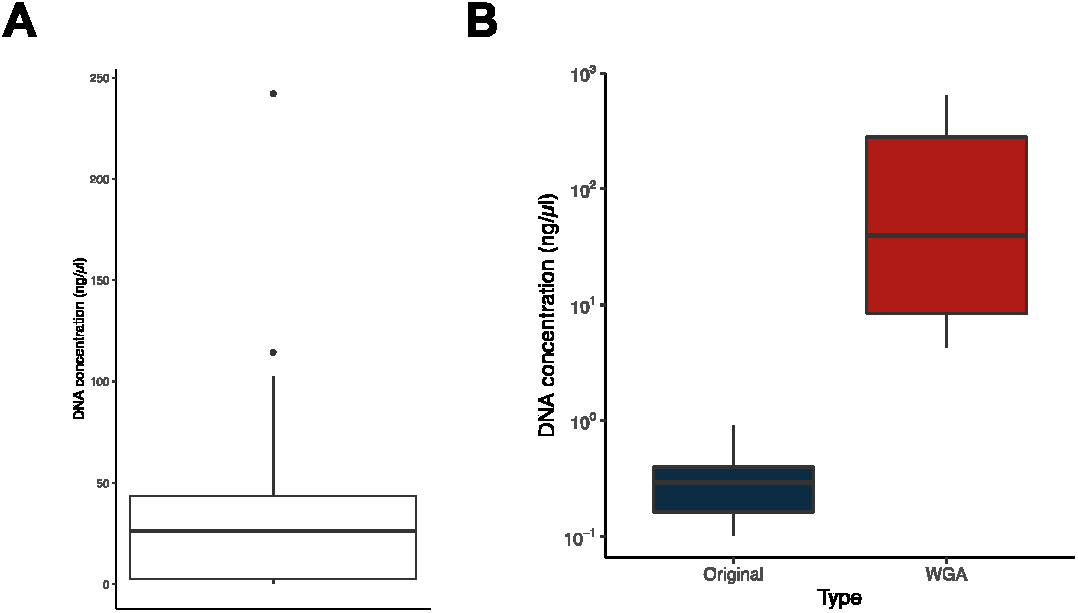
Whole genome amplification (WGA). **(A)** Box plot showing the distribution of double-stranded (ds) DNA concentration in the whole cell extracts (datasets Batches 2 and 3). **(D)** Box plots showing the distribution of dsDNA in original samples and in the same samples subjected to WGA.

To recover the remaining samples (n= 52), which represented 17.2% of the total samples, and 62.7% of the samples from the 2019 – 2021 period, we took advantage of Phi29-dependent whole genome amplification (WGA). As shown in Figure 2B, we have successfully increased the dsDNA concentration in our samples via WGA and hence proceeded with the WGS of these DNA-enriched samples.

### Recovery of mycobacterial reads from heavily contaminated DNA samples

From a total of 302, we successfully sequenced 270 samples (89.4% of the initial set), of which 244 were original (*i.e.*, not amplified) and 26 samples were subjected to WGA [[13] and this study]. The remaining either failed the initial DNA quality control coupled to the sequencing service or failed at library construction, due to DNA degradation or low amount of library DNA obtained. These 302 samples were sequenced in three batches: Batch 1 is that of Reis *et al.* [13], Batch 2 contained the remaining samples up to 2018, and Batch 3 contained the samples from 2019 onwards.

For each of the successfully sequenced samples, we checked the quality of the reads with FastQC [21] and plotted the categorical classification by type of sample – *i.e.*, those from Batch 1 [13], those from Batch 2, and those of Batch 3 divided in original and WGA (Figure 3A). The most prominent differences concern the “Per base sequence content”, “Per sequence GC content”, “Sequence Duplication Levels”, “Overrepresented sequences”, and “Sequence Length Distribution”.

**Figure 3.**
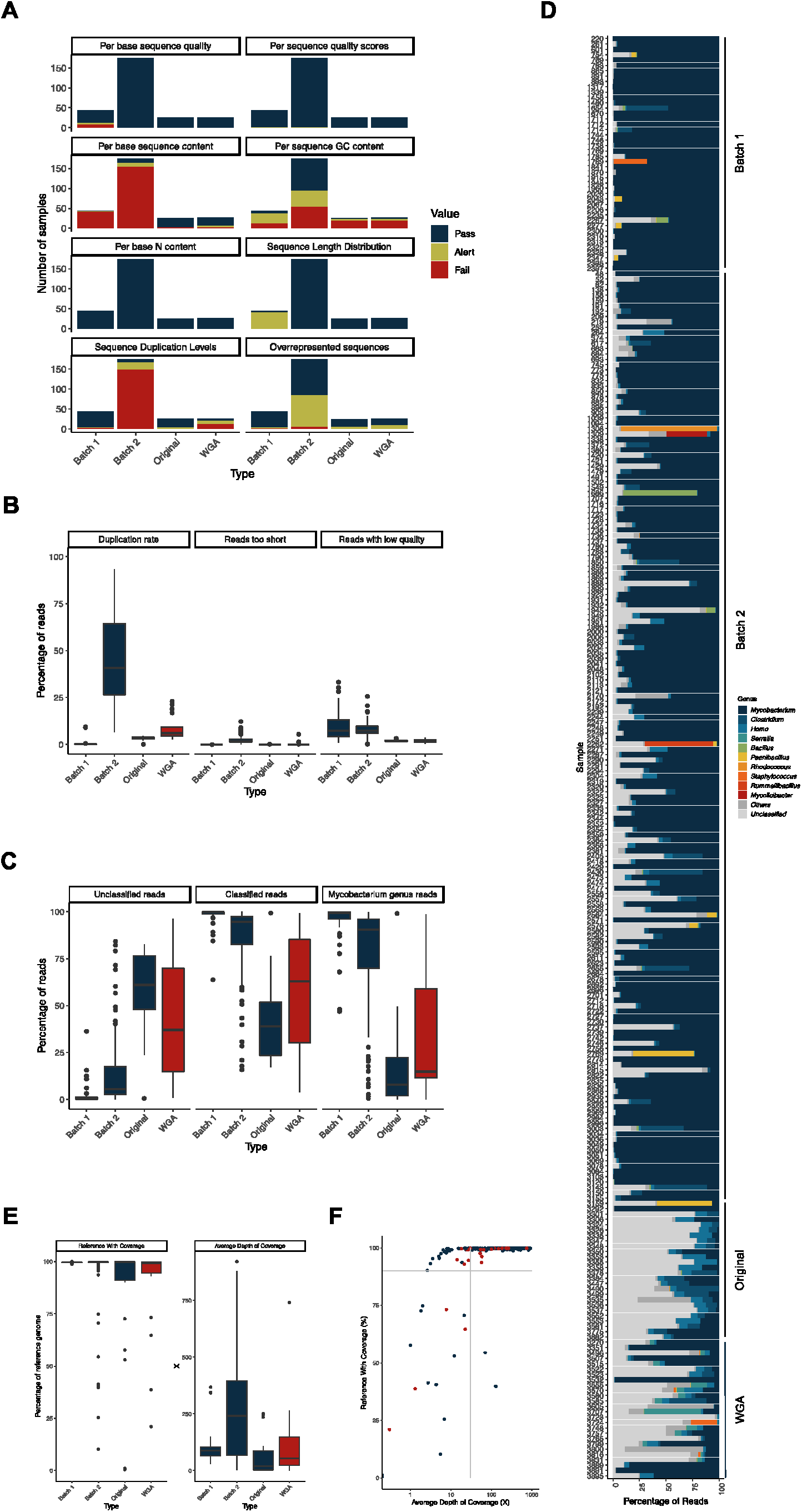
Whole genome sequencing. **(A)** Comparison of the sequencing reads quality parameters assessed by FastQC [21], divided by datasets Reis *et al*. [13] (Batch 1), Batch 2 and Batch 3 (original and WGA). FastQC classified the samples into three qualitative categories: Pass, Alert, and Fail. **(B)** Box plots of the distribution of the parameters analyzed by FastP [22] in the samples from the same dataset of panel A. The determined values of duplication rate, reads too short, and reads with low quality are percentages of the total of reads. **(C)** Box plots of the distribution of the Kraken2 [24] determined percentages of unclassified, classified, and mycobacterial reads in each sample. The samples were grouped by dataset, as in panel A. **(D)** Percentage of reads per sample classified in each of the genera indicated on the right. The “Unclassified” category comprises all the reads Kraken2 [24] was not able to classify and the “Other” category comprises the total of reads classified to other genera or not classified at the genus level. Represented are the ten most abundant genera in the full sample set. Datasets are indicated on the right. **(E)** Box plot of the distribution of the vSNP determined parameters “Average Depth of Coverage” and “Reference With Coverage”. **(F)** Distribution of the samples – Original and WGA – by the percentage of the reference with coverage and the average depth of coverage. The grey lines indicate the adopted quality thresholds of 90% reference with coverage and 30X average depth of coverage. All datasets are included here and those of Reis *et al*. [13] (Batch 1) and Batch 2 were colored blue because they have not been subjected to WGA.

As for the “Per base sequence content” category, most of the samples from Batches 1 and 2 raised an Alert or Fail (95.5% and 93.1%, respectively), indicating a non-uniform distribution of bases along the sequence. This is most likely related to the sequencing library construction kits used, as they differ in each case, and it is known that some kits generate this imbalance in base distribution. As for the “Per sequence GC content” category, 34.4% of samples fail in this criterium. This classification occurs because of the presence of sequences with very distinct GC content, being an indication that DNA of several different organisms was present in the samples. This does not seem to be a consequence of WGA, as the proportion of samples that fail this criterium is similar between original and WGA samples in the same period (2019 to 2021; 69.2% in the original and 64.5% in the WGA samples). As for the “Sequence Duplication Levels” category, we observed a high number of “Fail” occurrences in the Batch 2 dataset (84.5%) which is most likely related to the specificities of the sequencing, as the library kit used (TruSeq kit) is known to generate high levels of sequence duplication. The results of the FastP analysis make this aspect of increased duplication also evident, with much higher duplication rates observed in the samples of Batch 2 and those subjected to WGA (Figure 3B). Additionally, an increase in the levels of sequence duplication in the WGA samples when compared with the original samples of the same period was registered, in agreement with previous reports [36]. As for the “Overrepresented Sequences” category, we again observe a high level of “Alert” classification in Batch 2 dataset, again due to the specificities of the sequencing. Finally, as for the “Sequence Length Distribution”, a higher number of alerts comes in Batch 1 dataset, reflecting a distinct sequencing technology that generated longer reads (250 bp vs. 150 bp in the other two datasets) [13].

As mentioned above, FastQC results suggest that more than one genome (and hence, organism) was present in several samples. Taxonomical classification of the reads made this particularly evident. Especially in the most recent samples (Original and WGA), a high percentage of reads was not classified by Kraken2 Standard library and the percentage of reads classified as belonging to organisms of the genus *Mycobacterium* was less than 50% in over half of the samples (Figure 3C). Looking to each sample individually (Figure 3D), it becomes evident the presence of reads classified as belonging to other organisms and high percentages of reads that could not be classified in the most recent samples. The genus of the most frequent contaminants in all samples were *Homo* (human DNA), most likely from cross-contamination in sample manipulation, and other known animal pathogens or regular members of animal’s microbiota (*i.e.*, *Clostridium*, *Serratia*, *Bacillus*, *Paenibacillus*, *Rhodococcus*, *Staphylococcus*, *Rummeliibacillus*, and *Mycolicibacter*) [37–45]. One aspect that stands out is that the percentage of contamination with Gram-negative pathogens is higher in the WGA samples (average of 6.0%) than in the original (non-amplified) samples (average of 0.3%) from 2019 to 2021. Since each sample is different from one another, we cannot exclude that this result might, by chance, be related to these particular samples, but the hypothesis that WGA has generated a bias towards these bacteria cannot be excluded.

To overcome this issue, we have filtered the mycobacterial reads before proceeding with the alignment to the *M. bovis* AF2122/97 reference genome. This filtering step was performed whenever the percentage of *Mycobacterium* reads in the sample was below 70%. As shown in Figure 3E, the obtained values of average depth of coverage were higher in the Batch 2 dataset and on the WGA samples, especially when compared to the Original samples (Batch 3). Conversely, the coverage of reference genome was lower in the WGA sample set than in the Original one (Figure 3E). This is most likely due to the low percentage of mycobacterial reads in several of the WGA samples (Figure 3D) along with higher duplication rates in these samples (Figure 3B).

To include these samples in our ongoing whole genome-based epidemiological surveillance, we applied quality thresholds to our alignments to allow for robust SNP calling: at least 90% coverage of the reference genome and at least 30X depth of coverage. Based on these criteria, 201 samples (74.4% of the sequenced samples) were deemed suitable for inclusion in our ongoing epidemiological analysis (Figure 3F).

Note that other non-tuberculous mycobacteria can be isolated from TB-compatible lesions [46], and consequently generate *Mycobacterium* genus reads. Although our starting material (archived DNA obtained from cultured strains) had already been tested by molecular methods for the presence of *M. bovis* DNA (see Materials and Methods for details), we assessed the presence of mycobacterial reads other than from *M. bovis* by analysing the reads that did not align with its reference genome. We have observed that the percentage of unaligned reads was very low, both in the samples in which the reads were taxonomically filtered due to high levels of contamination (*i.e.*, above 30%) or those that were not (Figure S1A). As expected, the percentage of mycobacterial reads in those unaligned was higher in the filtered samples (Figure S1B) but they were assigned at very low percentages to several other mycobacterial species (Figure S1C), sometimes even with only one read being assigned to a given species. Altogether, these data do not support contamination of our samples with other non-tuberculous mycobacteria.

We have also evaluated how our sequencing metrics of quality and contamination varied with the origin of the sample, as those of wildlife are collected in the field and may be more prone to contamination. However, no significant differences were observed between samples of different host origins in neither of the evaluated metrics (Figure S2), which indicates that the results and conclusions we obtained can be generalized to all sample origins (*i.e.,* host species).

### Genome separation in cases of mixed infections

While aligning the filtered reads to the reference genome with vSNP, multiple *M*. *bovis* isolates were detected in five samples, *i.e.*, five samples in which defining SNPs had ambiguous calls. To recover samples 219, 1790, 1923, 2110, and 2244, SplitStrains was used [28]. The mixed *M*. *bovis* isolates were separated into ten strains, with the major strains occurring in proportions between 0.62 and 0.95 (Table 1). Of these ten separated samples, seven samples passed the quality thresholds (Table 1) and were included in the whole genome epidemiological surveillance dataset. Hence, this approach allowed us to recover 70% of otherwise unusable samples.

**Table 1.**
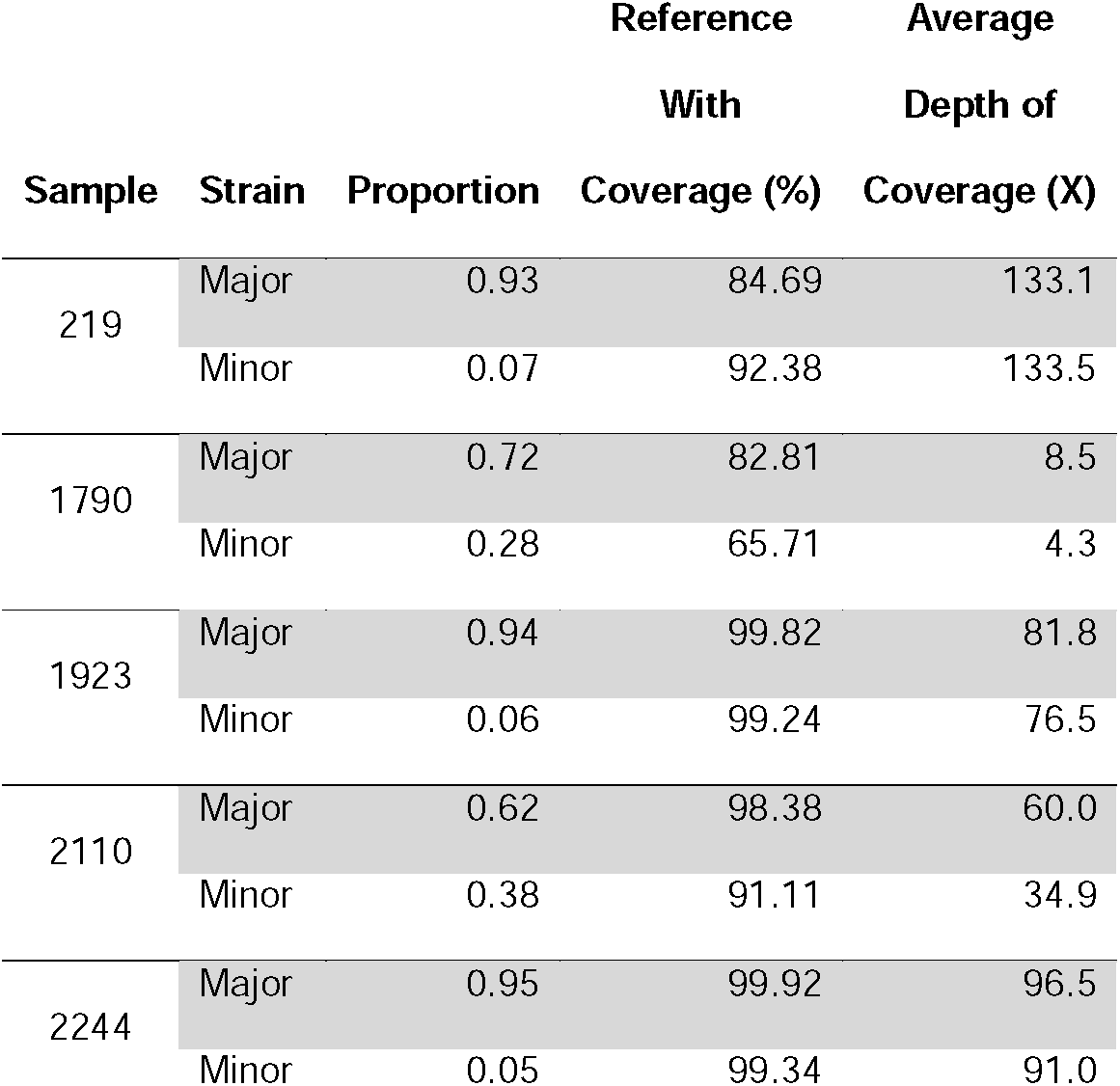
Mixed isolates separation. Analysis was performed with SplitStrains [28] and vSNP.

### Recovery of fragmented assemblies

Not all relevant analyses (*e.g.* comparative and functional genomics) can be done under a map-to-reference approach and, in such cases, it becomes necessary to use the sequencing reads to generate a genome assembly. Hence, we generated genome assemblies of the 213 samples that passed the map-to-reference sequencing depth and genome coverage quality thresholds (30X and 90%, respectively). We assembled the reads with Unicycler [47] and analyzed the quality of the resulting assemblies with Quast [35] (Table S2). We looked at the data the same way as in the map-to-reference data, and saw that the quality of the assemblies is much lower in the samples that were subjected to WGA (n= 12) (Figure 4). In fact, not only was the number of contigs generated higher (Figure 4A), but also the number of bases covered in the assembly was lower (Figure 4B). In addition, the assembler was unable to generate large contigs (Figure 4C), and the percentage of reference genome covered by the assemblies was much lower (Figure 4E), when compared with the samples not subjected to WGA (*i.e.*, those of Batches 1 and 2, and the Original samples of Batch 3).

**Figure 4.**
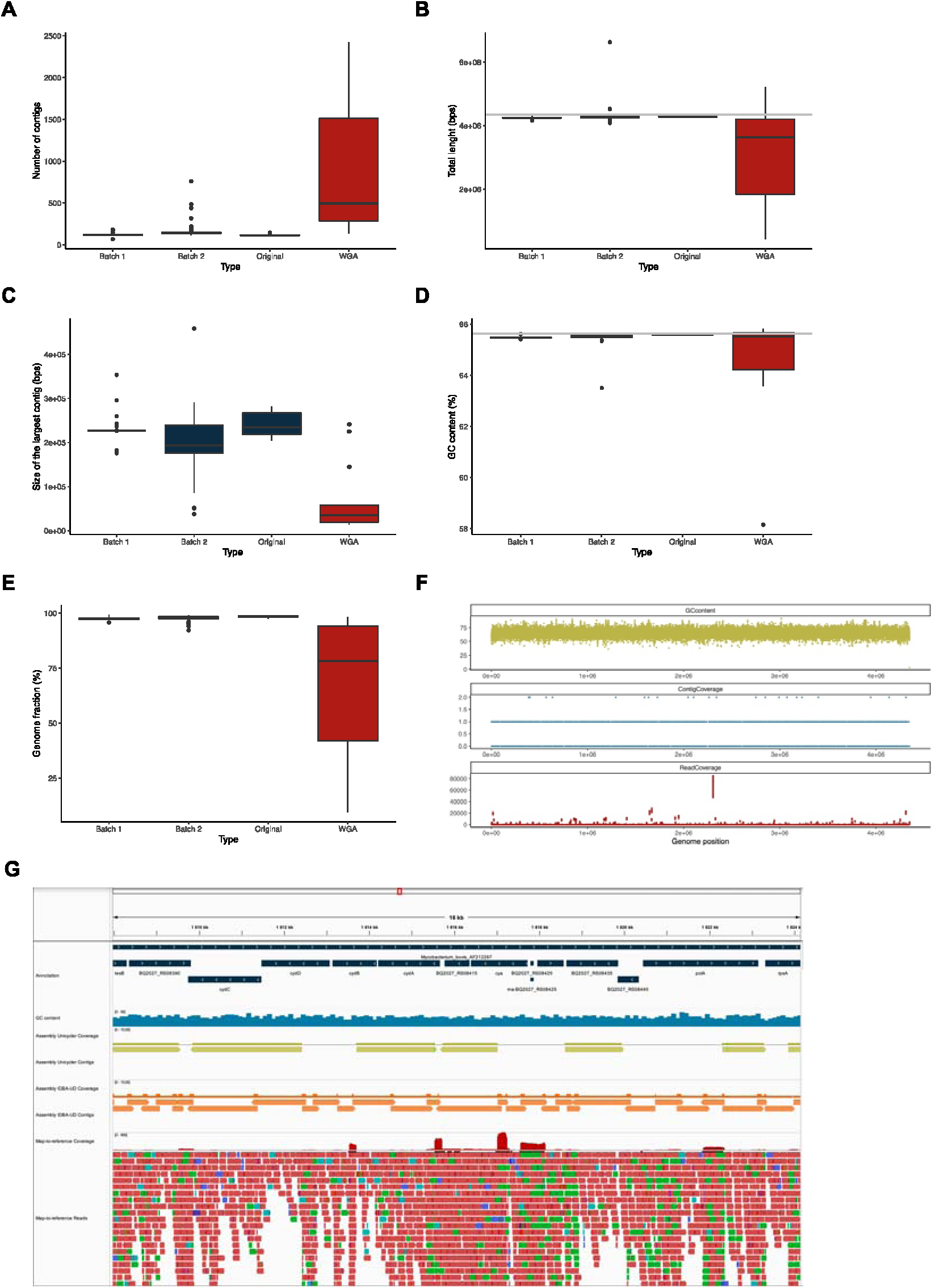
*De novo* assembly. **(A-E)** Box plots of the distributions of the number of contigs **(A)**, total length of the sum of all contigs **(B)**, size of the largest contig **(C)**, GC content **(D)**, and genome fraction **(E)** of the *de novo* assemblies generated by Unicycler and assessed by Quast, divided by the original datasets of Reis *et al*. [13] (Batch 1), Batch 2, and Batch 3 (original and WGA). **(F)** Variation of GC content (calculated in 100 bp windows), contig coverage and read coverage in the reference genome of *M. bovis* AF2122/97 of sample 3270. **(G)** Snippet of a genome region showing the genome annotation (dark blue), the GC content (blue), the coverage by contigs generated by Unicycler (green) and IDBA-UD (orange), and the coverage of the reads (red) in sample 3270.

We also noticed that the assemblies of the samples subjected to WGA had a much lower GC content (Figure 4D), which led us to hypothesize that the low quality of the assemblies was due to the WGA bias towards low GC content sequences, which would generate a bias in the amplification reducing the quality of the assemblies. However, such hypothesis would not be in agreement with the high genome coverages obtained with the map-to-reference approach (see above) and was also not in agreement with what was seen by plotting the reference genome GC content versus the contig coverage (see Figure 4F for the full extent of the genome and Figure 4G for an insert into a 16 kb region) or previous descriptions [48]. Despite refuting our hypothesis, this analysis highlighted the lack of contigs in areas where the read coverage was very high.

To further understand this aspect, we analized 25 sequenced samples that were previously subjected to WGA and 26 randomly chosen samples selected from amongst those that were not, concerning map-to-reference mapping quality, depth of coverage and SNP density (Figure S3). This analysis made particularly evident the uneven coverage throughout the entire *M. bovis* chromosome, while no obvious differences were observed regarding mapping quality or SNP density, with both sample types generating very similar results. This uneven sequencing depth has already been noted in samples subjected to WGA (as were those of single cell sequencing [48]) and prompted us to use an assembler specifically developed to deal with such samples – *i.e.*, IDBA-UD [32]. By using this assembler and posterior contig fusion, we were able to generate four additional quality assemblies (Table S3) and rescue 36.4% of the WGA assemblies.

## Discussion

In this work, we explored different strategies to rescue DNA samples obtained by routine approaches performed by national reference laboratories for animal TB. Obtaining DNA in enough quantity and quality for WGS can be challenging when interconnecting the routine culture and PCR-based schemes with WGS-based methods. It is therefore important to have available a set of alternative approaches to recover samples that suffer from the most commonly found limitations, *e.g.*, low amount of DNA, high rates of contamination, and mixed infections (*i.e.*, infections with different strains of the same pathogen). Why is it so important to recover as many *M. bovis* samples as possible? Because the loss of an isolate’s genome represents the loss of a link in the reconstruction of transmission chains, hampering the interpretation of the biological and epidemiological data as a whole.

Following routine procedures, the amount of DNA recovered from the samples processed in the automatic growth detection system was low, hindering direct WGS in 17.8% of total samples (which correspondend to 62.7% of the samples recovered directly from the automatic growth detection system cultures) (Table S4). To overcome this issue, we applied WGA and successfully amplified all samples, with an increase in the amount of DNA from 9 to 2025 fold. This approach enabled the recovery of all samples for the sequencing workflow, *i.e.*, we obtained at least 100 ng of total DNA for all 56 samples.

Another recurrent issue was the contamination with DNA from organisms other than *M. bovis*, particularly evident in the most recent samples (*i.e.*, from sample number 3270 onwards) (Table S4). The decontamination is made with 4% sodium hydroxide, that kills more sensitive cells but does not necessarily eliminate the DNA present in the sample, because it might still be protected by cellular structures (*e.g.* endospores) or even if free, it may suffer denaturation but not complete degradation [49]. The culturing step should increase the amount of mycobacterial DNA by allowing mycobacterial multiplication, but if the cultures are processed immediately after being positive for growth (as assessed by fluorescence, which is a very sensitive indicator) that enrichment might not be sufficient. One methodological change that might minimize this issue would consist of extending the incubation time of the culture tubes to allow additional growth of mycobacteria.

The bacterial genera that appear in these samples, all of which have been associated with animal infections or the animal’s regular microbiota [37–45], suggest that the contamination source is related with the sampled animals and not with posterior contamination introduced by handling of the samples. Of course, if such organisms are present in a high amount in the sampled tissues, and since the decontamination procedure does not eliminate the DNA (as discussed above), they will generate WGS reads.

Another hypothesis that could explain this observation is that the organisms we find are resistant to the decontamination process and the antibiotics present in the growth medium. The antibiotics included in the PANTA supplement target fungi (amphotericin), Gram-negative bacteria (polymyxin B, nalidixic acid, trimethoprim, and azlocillin), and a small group of Gram-positive bacteria (trimethoprim; *Staphylococcus* and *Streptococcus*). Hence, if the contamination we found would have come from growth of the microorganisms in the MGIT tube, they would have to resist the decontamination process along with being resistant to the antibiotics present in the medium. *Clostridium* spp., the most common contaminants, are strict anaerobes, which makes the growth in the MGIT tube unlikely, but their spores could be resistant to decontamination and hence their DNA detected. The same is true for other spore forming organisms that were also detected (*i.e.*, *Bacillus* spp., *Paenibacillus* spp., and *Rummellibacillus* spp.) or other actinobacteria (*i.e.*, *Rhodococcus* spp. and *Mycolicibacter* spp.), whose more robust cell envelope might make them more resistant to the decontamination protocol. Finally, the carbon sources available in the Middlebrook medium (supplemented with OADC) are not primary carbon sources readily used by non-mycobacterial organisms. Altogether, growth of contaminants seems unlikely in these conditions.

We have attempted to overcome this high contamination issue by filtering out the mycobacterial WGS reads. In almost a quarter of all sequenced samples, the fraction of mycobacterial reads was low and the alignment to the reference genome did not have sufficient quality to be included in the subsequent whole-genome based epidemiological analysis. One possible way to overcome this issue is increasing the sequencing depth (*i.e.* the number of reads generated with WGS) as a way to increase the number of mycobacterial reads. Another approach might rely on whole genome enrichment, which was already successfully applied to enrich *Mycobacterium tuberculosis* in sputum samples [50].

Another challenge was the presence of more than one *M. bovis* strain in the same sample, as defined by the presence of ambiguous defining SNPs. This phenomenon might occur if two *M. bovis* strains independently infect the same animal host or if clonal evolution occurs inside the host during the typical TB chronic infection phenotype [51]. Strain separation, although effective, leads to an expected reduction in the percentage of genome coverage and average sequencing depth, given that the reads containing the SNPs are separated between the two resulting strains – *i.e.*, major and minor strains. Despite this additional limitation, seven out of the ten resulting strains could be rescued and included in the ongoing genomic epidemiology studies.

While generating *de novo* genome assemblies, we encountered a limitation related to WGA, *i.e.*, in the samples subjected to this amplification process, the assemblies had very low continuity. Our analysis correlated the areas in which a contig could not be generated to regions of extremely high read coverage. Given that most of the *de novo* assemblers identify regions with above average coverage as repetitive regions and break the contigs on those regions for not being able to unequivocally assign a read to a given repeat region, we were prompted to explore assemblers especially designed to generate *de novo* assemblies from data with uneven coverage. Our approach allowed us to recover four out of the 11 WGA samples that failed in the regular *de novo* assembly, but this must be taken into account when considering the use of WGA to rescue samples with low amount of DNA, as it is a strategy that performs better when the data is to be used for map-to-reference approaches, than for *de novo* assemblies.

A subset of the samples analyzed in this study had been previously used in published studies. Specifically, the map-to-reference alignments were used for phylogenomic analysis of the *M. bovis* population circulating between wildlife and livestock in Portugal [13], while *de novo* assemblies were used to support pangenome analysis [52]. The remaining samples (83.7%) have been incorporated into our ongoing studies encompassing both phylogenomics and phylodynamics (relying on the generated map-to-reference alignments), as well as the expansion of the pangenome analysis (using the *de novo* assemblies). These recovery strategies have not only prevented the loss of several genomes but have also permitted the coherent integration of these genomes into the *M. bovis* population phylogenetic trees for Portugal. This integration occurred without raising any suspicions of potential artifacts introduced by our recovery methods. In other words, these genomes distributed across various branches of the phylogenetic trees without any noticeable clustering (Pereira et al., unpublished). This outcome was entirely expected due to the lack of any disparity in SNP density across genomes subjected to whole genome amplification (WGA) compared to those that were not. Regarding the strains recuperated through read splitting, their positioning in the tree indicated that they are unrelated, suggesting instances of mixed infections rather than evolution within a single host (Pereira et al., unpublished). Consequently, when these recovered genomes are applied in subsequent analyses, their behavior aligns with that of genomes that did not require the implementation of recovery strategies. This reinforces the validity of the recovery strategies employed in this study and sustains the inclusion of the resulting genomes in downstream epidemiological and evolutionary analyses.

In conclusion, when adding to the routine culture and PCR-based detection of *M*. *bovis*, a series of procedures leading to the obtention of a complete genome sequence, one might expect additional challenges. These challenges take form in the contamination of the sample, the low amount of available DNA, and the potential presence of multiple *M. bovis* strains in the same sample (Table S4). Despite undesirable, these issues can be overcome. Here we showed one way to overcome these challenges, but others could certainly be explored with the necessary adjustments to the particularities of the samples and resources available. We were able to recover 100% of the samples with a low amount of total DNA through WGA, 60.8% of the contaminated samples through read filtering, and 70% of the mixed strains through strain separation. Overall, these approaches allowed the recovery of 62 samples that would have been otherwise unusable for epidemiological purposes.

Thus, the direct coupling of genomics to *M. bovis* standard diagnosis is possible provided that specific laboratory and computational strategies are applied to circumvent the limitations imposed. Modification of the diagnostic protocol and a dedicated pipeline integrating computational tools tackling different data challenges minimises biological data losses, optimizes available material resources, and greatly facilitates the coupling of diagnosis to genomic surveillance. In fact, results shown here also argue in favor of the inclusion of an additional step of culture of *M. bovis* in solid medium after the automatic detection whenever WGS is the final goal. The advantage of including this step, present in the conventional method but not in the automatic one, is particularly obvious in the amount and purity of the DNA that can be retrieved. This would facilitate the downstream computational analysis and the likelihood of obtaining useful sequencing data for genomic epidemiology purposes.

Here, we have applied these rescue strategies to our particular problem of recovering *M. bovis* genomes to answer animal TB related epidemiological questions. However, the value of the implementation of these rescue strategies is not limited to our particular model system and has a broader application. In fact, they can be applied to any other sample type in which similar problems might arise, as is the case of blood cultures for diagnosis of bloodstream infections, where the percentage of reads belonging to the pathogen is low [53], impairing the capability of predicting antibiotherapy susceptibility from WGS data [54], amongst other limitations [55].

## Acknowledgements and Funding

We thank Rogério Tenreiro and Rute Fonseca for technical suggestions. We acknowledge funding from Fundação para a Ciência e a Tecnologia, IP (FCT)/MCTES through national funds (PIDDAC) and co-funding by the European Regional Development Fund (FEDER) of the European Union, through the Lisbon Regional Operational Program and the Competitiveness and Internationalization Operational Program for Portugal 2020 or other programs that may succeed it in the scope of project “Colossus: Control Of tubercuLOsiS at the wildlife/livestock interface uSing innovative natUre-based Solutions” (references PTDC/CVT-CVT/29783/2017, LISBOA-01-0145-FEDER-029783, POCI-01-0145- FEDER-029783). Strategic funding from FCT to cE3c and BioISI Research Units (UIDB/00329/2020 and UIDB/04046/2020) and to the associate lab CHANGE (LA/P/0121/2020) are gratefully acknowledged. ACP was funded by FCT (SFRH/BD/136557/2018).

## Data Accessibility

The sequence data included in this work are deposited under Bioproject accession numbers PRJNA682618 and PRJNA946560 at a public domain server in National Centre for Biotechnology Information (NCBI) SRA database.

## Author Contributions

DP and MVC designed the study. AB provided the DNA samples. DP, ACP and GT performed the computational analyses. DP, ACP, GT and MVC analyzed the output data. DP performed visualizations. DP and MVC wrote the manuscript with contributions from ACP and GT.

## Supplemental information

**Table S1. Full dataset.** The data of all samples concerning the source of the sample, amount of DNA, read quality control and map-to-reference metrics are given in table format.

**Table S2. *De novo* assemblies.** The quality control data for all generated assemblies is given in table format.

**Table S3. *De novo* assemblies of WGA data.** The quality metrics of the *de novo* assemblies of WGA samples using Unicycler, IDBA-UD and a combination of IDBA-UD and Samtools is shown for comparison.

**Table S4. Summary Table.** The number of samples that passed each quality criterium is summarized.

**Figure S1.**
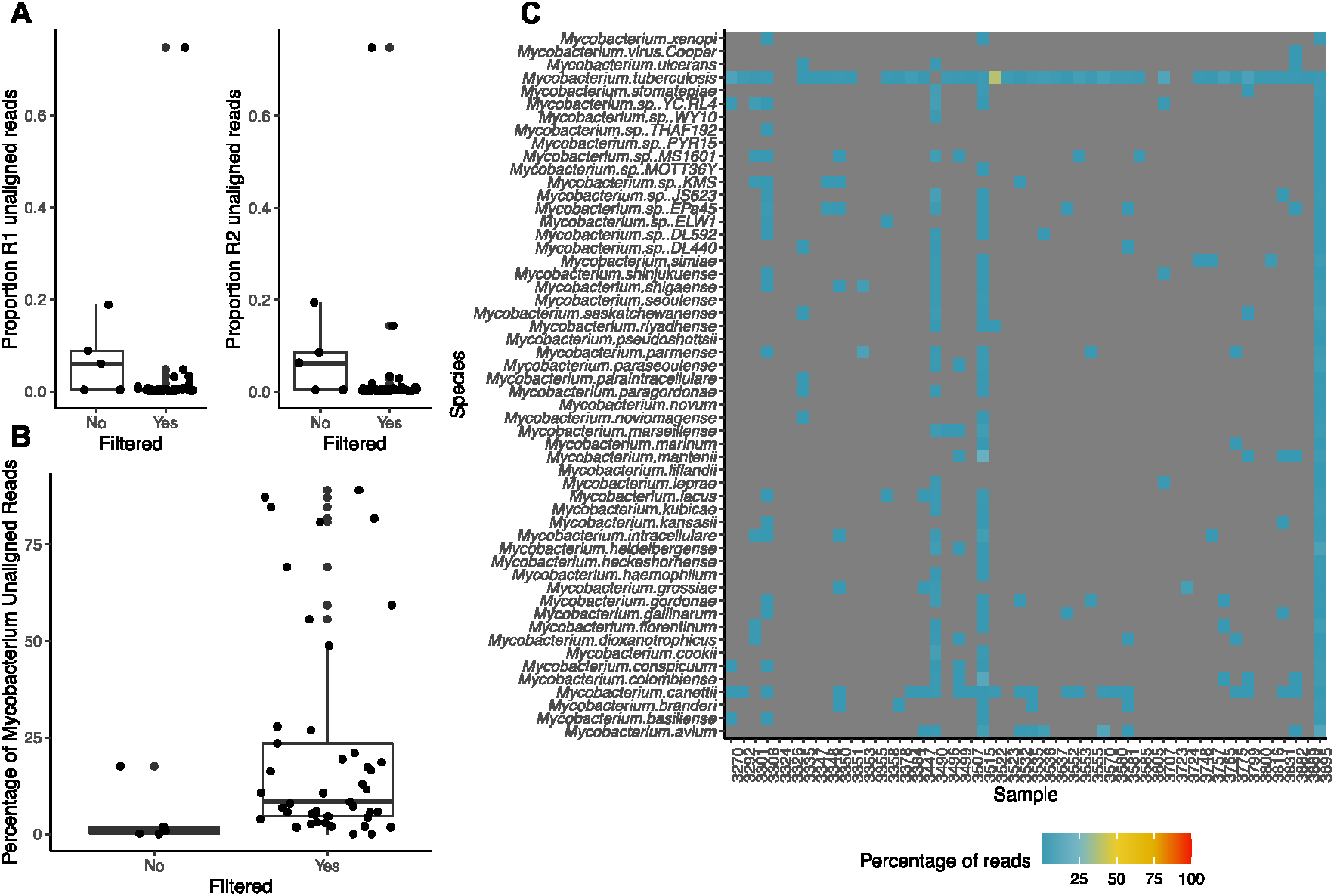
Contamination with non-tuberculous mycobacteria. **(A)** The distribution of the proportion of unaligned reads in both directions (R1 and R2) for Batch 3 samples that either were filtered to keep only *Mycobacterium* genus reads or not. **(B)** Percentage of those unaligned reads that once reclassified by Kraken were again classified as *Mycobacterium* reads. **(C)** Percentage of those *Mycobacterium* genus unaligned reads that were assigned to a given species. Grey means that no unaligned reads of that sample were assigned to that species.

**Figure S2:**
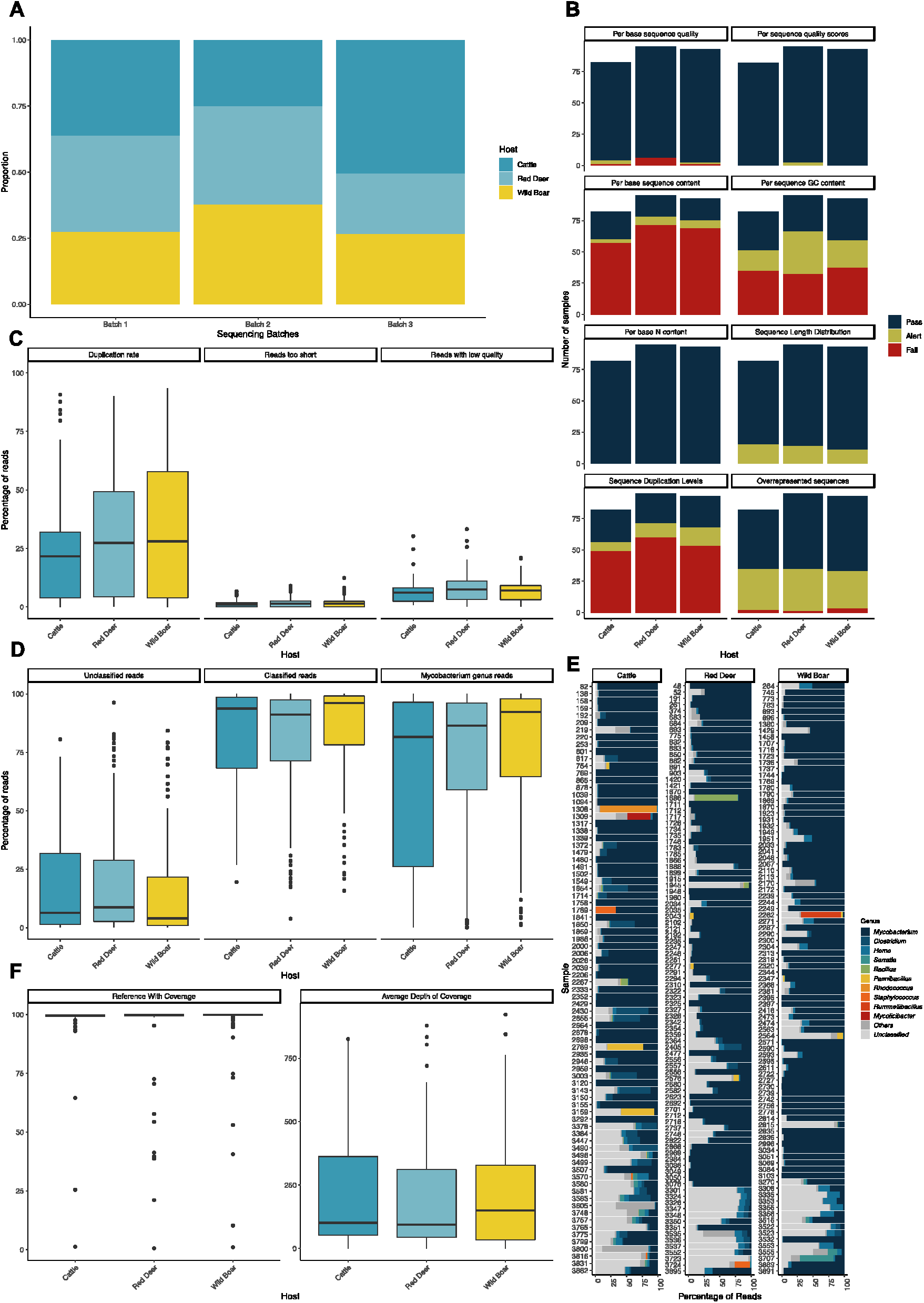
Sequencing metrics by the origin of the sample. **(A)** Distribution of host species in each sequencing batch. **(B)** Comparison of the sequencing reads quality parameters assessed by FastQC (Andrews, 2010), divided by the origin of the sample (cattle, red deer, or wild boar). FastQC classified the samples into three qualitative categories: Pass, Alert, and Fail. **(C)** Box plots of the distribution of the parameters analyzed by FastP (Chen et al., 2018) in the samples from the same dataset of panel A. The determined values of duplication rate, reads too short, and reads with low quality are percentages of the total of reads. **(D)** Box plots of the distribution of the Kraken2 (Wood et al., 2019) determined percentages of unclassified, classified, and mycobacterial reads in each sample. The samples were grouped by dataset, as in panel A. **(E)** Percentage of reads per sample classified in each of the genera indicated on the right. The “Unclassified” category comprises all the reads Kraken2 (Wood et al., 2019) was not able to classify and the “Other” category comprises the total of reads classified to other genera or not classified at the genus level. Represented are the ten most abundant genera in the full sample set. Sample origins are indicated on top. **(F)** Box plot of the distribution of the vSNP determined parameters “Average Depth of Coverage” and “Reference With Coverage”.

**Figure S3.**
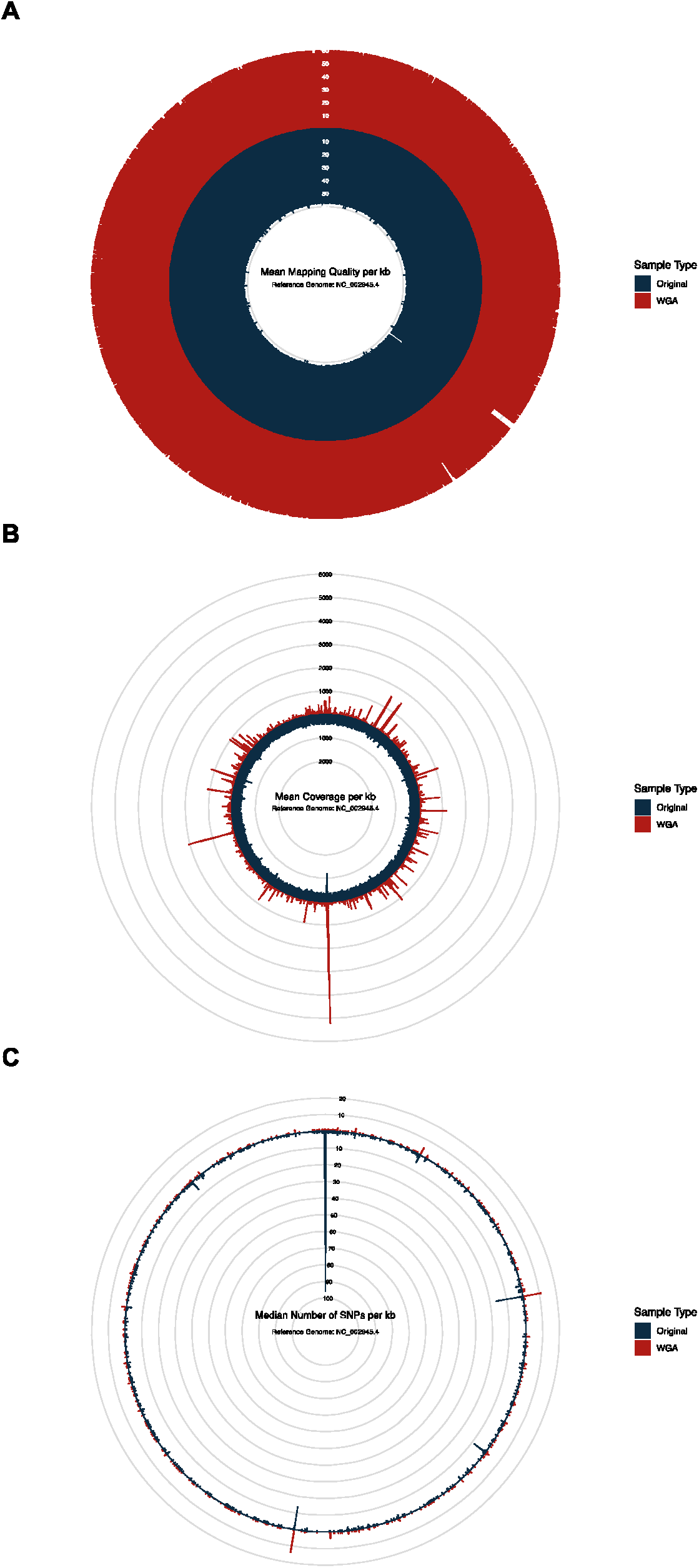
Comparison of mapping statistics between samples subjected or not to WGA. Twenty-five sequenced samples subjected to WGA and 26 randomly selected samples not subjected to WGA were analyzed. The circular plots show how mapping quality **(A)**, coverage **(B)** and SNP density **(C)** per kb change across the genome (on the circular axis). The data is color coded accordingly to the sample type (WGA or original).

